# A minimal transcriptomic signature predicts intravascular tumor extension in renal cell carcinoma

**DOI:** 10.64898/2026.03.05.709839

**Authors:** Christopher A. Mao, Ramiro Ramirez, Hanzhang Wang, Wasim H. Chowdhury, Dharam Kaushik, Ronald Rodriguez

## Abstract

Renal cell carcinoma (RCC) with venous tumor thrombus, termed renal intravascular tumor extension (RITE), is associated with aggressive behavior and poor clinical outcomes. Yet, its underlying molecular determinants remain incompletely defined. We analyzed RNA sequencing data from three independent RCC cohorts comprising 721 samples. Two cohorts included matched samples of index tumor, tumor thrombus, and normal adjacent kidney tissue. Analyses integrated dimensionality reduction, differential gene expression, interpretable machine learning, and gene ontology approaches. Principal component analysis revealed that only these two cohorts exhibited a coherent RITE-associated transcriptional structure. Their sequencing depth was sufficient to delineate 6,317 differentially expressed genes that distinguish RITE from non-RITE tumors. SHAP-based feature attribution across logistic regression, random forest, and XGBoost yielded a robust 29-gene consensus signature, which was further distilled into a compact 13-gene panel that preserved maximal classification performance. These genes converged on biological themes, including loss of distal epithelial identity, dysregulation of ion transport pathways, and consistent enrichment of mitochondrial processes such as oxidative phosphorylation. Together, these findings define a newly discovered and uniquely refined molecular signature of venous tumor extension in RCC and highlight mechanistically relevant pathways that may inform biomarker development and future translational strategies for predicting or mitigating RITE progression.

## Introduction

Renal cell carcinoma (RCC) originates from renal epithelium and accounts for more than 90% of cancers in the kidney^1^. A distinctive and clinically aggressive feature of RCC is its propensity to extend directly into the venous circulation, forming a tumor thrombus within the renal vein or inferior vena cava^1–3^. This phenomenon, referred to as renal intravascular tumor extension (RITE)^4^, occurs in approximately 15% of RCC patients and represents one of the most surgically challenging manifestations of the disease^4,5^.

Clinically, RITE is associated with worse outcomes, including higher rates of synchronous metastasis, increased perioperative morbidity, and life-threatening complications such as pulmonary embolism and cardiac involvement^3,5–8^. In the absence of definitive treatment, patients with RITE exhibit poorer prognosis, with reported median survival as low as five months and a one-year disease-specific survival rate of approximately 29%^9^. Although advances in surgical techniques, including complex thrombectomy and cardiopulmonary bypass, have improved survival for select patients, these procedures carry significant operative risk, and perioperative mortality remains nontrivial^3,10–12^. Moreover, while neoadjuvant systemic therapies, including tyrosine kinase inhibitors and immune checkpoint inhibitors, have been explored in this setting, they have demonstrated limited benefit in reducing tumor thrombus burden. In some cases, these therapies may induce local inflammatory reactions that increase the technical complexity of surgical excision^3^.

Given these challenges, early identification of tumors at risk for venous invasion along with a mechanistic understanding of RITE are critical unmet needs in RCC management. The molecular programs that enable RCC cells to invade and survive within the venous microenvironment remain incompletely understood. Prior studies have implicated pathways related to angiogenesis, hypoxia, epithelial-mesenchymal transition, and immune modulation in aggressive RCC phenotypes^4,13,14^.

A major barrier to progress in this area is the lack of explicit RITE annotations in widely used public datasets. Large-scale transcriptomic resources such as The Cancer Genome Atlas Kidney Renal Clear Cell Carcinoma (TCGA-KIRC)^15^ cohort do not systematically label venous invasion or tumor thrombus status, limiting their utility for studying RITE-specific biology. Consequently, transcriptomic signatures uniquely associated with venous extension may be diluted or obscured when invasive and non-invasive tumors are analyzed together.

To address these gaps, we performed transcriptomic analyses that explicitly compare tumor thrombus, tumor index, and normal renal tissues in well-characterized RCC cohorts. Such analyses elucidate deregulated transcriptional homeostasis associated with RITE formation, identify molecular features that distinguish venous-invasive tumors from non-RITE counterparts, and uncover biologically meaningful targets for risk stratification and therapeutic development. To the authors’ knowledge, no published studies have systematically investigated RITE-associated transcriptomic dysregulation using machine learning frameworks combined with explainable artificial intelligence techniques to derive powerful yet interpretable biological insights.

In this study, we analyze three transcriptomic datasets—Rodriguez, Wang^16^, and TCGA-KIRC—to evaluate their suitability for RITE-focused analyses, identify differentially expressed genes distinguishing RITE from non-RITE tumors, and apply explainable machine learning approaches to derive concise, biologically-coherent gene signatures predictive of venous invasion.

## Methods

### Clinical sample collection and storage

This study was approved by the UTHealth San Antonio Institutional Review Board under protocol IRB #2020042HU. Written informed consent was obtained from all participants before any study procedures. All methods were carried out in accordance with relevant guidelines and regulations. Patients were eligible if they had RITE. Intraoperatively, intravascular tumor tissue, the non-venous portion of the tumor termed the index tumor, and adjacent normal kidney tissue were collected from patients undergoing nephrectomy with tumor thrombus resection. Immediately after surgical excision, a genitourinary pathologist macro-dissected all tissues and snap-froze them in either liquid nitrogen or a dry ice-ethanol slurry under RNase-free conditions. All tissue processing was completed within 2 hours of organ removal to preserve RNA integrity for downstream analyses.

### RNA extraction, library preparation, and sequencing

Tissues from the Rodriguez cohort were processed for bulk RNA sequencing (RNA-seq) as follows. Cells were harvested by washing twice with phosphate-buffered saline (PBS; ATCC^®^), followed by trypsinization and neutralization with complete media. Cell suspensions were centrifuged at 1,500 rpm for five minutes, washed once more with PBS, and pelleted. Total RNA was extracted by adding 500 µL of TRIzol^™^ Reagent (Sigma-Aldrich) according to the manufacturer’s protocol, which is based on acid guanidinium thiocyanate–phenol–chloroform extraction^17^. RNA quantity and purity were assessed by UV–visible spectrophotometry at 260/280 nm using a SpectraMax^®^ i3x Multi-Mode Microplate Reader (Molecular Devices^®^).

Library preparation was performed using approximately 500 ng of total RNA with the KAPA^®^ Stranded RNA-Seq Kit with RiboErase^®^ (Roche Sequencing Solutions). Ribosomal RNA was depleted, and the remaining RNA was fragmented using divalent cations under high-temperature magnesium conditions. Fragmented RNA was converted into double-stranded cDNA using random priming, followed by adapter ligation and PCR amplification to enrich the final sequencing libraries. Library size distribution and integrity were verified using an Agilent 2100 Bioanalyzer^®^ system (Agilent Technologies^®^), and concentrations were quantified with a Qubit^™^ Fluorometer using the Qubit^™^ dsDNA HS Assay Kit (Thermo Fisher Scientific^®^). Libraries were normalized to 20 nM and pooled for sequencing.

Sequencing was performed on an Illumina^®^ NextSeq^™^ 500 platform (Illumina^®^) using 75 bp paired-end reads, generating an average depth of approximately 35 million reads per sample. Raw binary base call files were demultiplexed to produce FASTQ files, which were imported into CLC Genomics Workbench^®^ version 21 (QIAGEN^®^) for initial processing. Reads underwent quality control and adapter trimming, followed by alignment to the *Homo sapiens* GRCh38/hg38 reference genome using default mapping parameters. Gene-level raw counts were then generated from the mapped reads.

### Data collection of external datasets

Because RITE is, by definition, confined to tumors with venous involvement, only TCGA tumors annotated as pathologic stage T3b or T3c (T3BC)—indicating tumor thrombus extension into the renal vein, inferior vena cava below the diaphragm, or inferior vena cava above the diaphragm—were considered RITE in downstream analyses. Tumors classified as T1 or T2, which lack venous invasion by definition, were retained exclusively as non-RITE comparators. Tumors annotated as T3a were excluded from analysis because this category encompasses heterogeneous patterns of local extension, including perinephric fat invasion, renal sinus fat invasion, and segmental renal vein involvement. These patterns cannot be reliably disaggregated within TCGA clinical annotations. As TCGA does not explicitly distinguish true intravascular tumor thrombus from other forms of T3a local extension, these tumors could not be confidently classified as RITE or non-RITE and were therefore removed to minimize biological ambiguity. Additionally, T4 tumors were excluded due to their advanced local invasion beyond Gerota’s fascia or into adjacent organs, representing a more advanced and biologically distinct disease state not directly comparable to RITE or non-RITE tumors.

These filtering criteria were applied to ensure that TCGA samples used in this study reflected a biologically coherent comparison aligned as closely as possible with true venous tumor extension.

In addition, whole-exome and transcriptome sequencing data from an independent RCC cohort published by Wang et al.^16^ were obtained from the National Center for Biotechnology Information (NCBI) Sequence Read Archive (SRA). Transcriptome sequencing data were accessed under accession codes `PRJNA596359` and `PRJNA596338`. Raw sequencing reads from these datasets were downloaded and processed locally to generate gene-level count matrices for downstream analyses.

### Computational environment

All computational analyses were performed in Google Colab (Google LLC), a cloud-based platform providing reproducible access to standardized hardware and software environments, using Python version 3.12.12. Analyses leveraged widely adopted, peer-reviewed libraries for statistical modeling, machine learning, and interpretability, including scikit-learn (v1.6.1) for model development and evaluation^18^, PyDESeq2 (v0.5.3) for differential expression analysis based on the DESeq2 framework^19,20^, XGBoost (v3.0.0) for gradient-boosted decision tree modeling^21^, SHAP (v0.50.0) for model interpretability using Shapley value-based feature attribution^22^, and statannotations (v0.7.2) for statistical comparison and visualization of model performance metrics.

### Principal component analysis (PCA)

PCA was used to evaluate global transcriptomic structure and cohort compatibility across datasets. The Rodriguez cohort and the external Wang cohort^16^ both include bulk RNA-seq profiles derived from surgically resected RCC specimens with explicit annotation of RITE, including non-vascular portions of the tumor (RITE-index), intravascular tumor thrombus (RITE-thrombus), and normal adjacent kidney tissue (RITE-NA) when available. In contrast, the TCGA-KIRC dataset comprises index tumor and normal adjacent tissue samples without explicit annotation of venous tumor thrombus.

Raw count matrices were normalized to transcripts per million (TPM). The TPM-normalized RNA-seq data were used for dimensionality reduction analyses. To reduce the influence of low-abundance noise, genes with an average raw count below one were removed independently within each dataset. Cross-cohort comparisons were performed using the intersection of genes shared among all three datasets, as well as separately within the Rodriguez/Wang^16^ and TCGA datasets to assess internal structure and cohort-specific effects. Because RNA-seq expression values span a broad dynamic range, data were transformed using log_2_(TPM + 1*e*^−6^) prior to PCA.

PCA was conducted using the scikit-learn library^18^ (version 1.6.1) in Python to evaluate clustering patterns based on overall gene expression profiles. For the Rodriguez and Wang^16^ cohorts, PCA assessed relationships among RITE-thrombus, RITE-index, non-RITE tumor, and normal adjacent samples. For TCGA, PCA was performed to examine the distribution of T1, T2, T3, and normal adjacent samples. A secondary mapping was applied in which T3 samples were labeled as RITE and T1/T2 samples as non-RITE, allowing an exploratory assessment of whether TCGA exhibited molecular structure consistent with RITE biology.

### Analyses of differentially expressed genes (DEGs)

Raw count matrices were analyzed using the PyDESeq2 package^19^ (version 0.5.3) in Python for normalization and differential expression testing.

Differential expression analyses compared RITE tumor samples with non-RITE clear cell renal cell carcinoma (ccRCC) samples to identify transcriptomic alterations associated with venous tumor extension. DEGs were determined in two settings. First, we analyzed RITE samples comprising the index tumor and tumor thrombus relative to non-RITE tumors in our cohort and in the dataset reported by Wang et al.^16^ Second, we performed the same comparison in the TCGA dataset.

Differential gene expression analysis was performed using raw RNA-seq counts. Because PyDESeq2 requires unnormalized counts and performs its own internal normalization^19^, TPM-normalized values were not used. To reduce noise, genes with an average raw count below one were removed independently within each dataset. Differential expression testing compared RITE tumors with non-RITE tumors in each cohort. For TCGA, tumor stage was used as a surrogate: T3BC tumors were classified as RITE^23^, and T1/T2 tumors as non-RITE. PyDESeq2 was applied using an adjusted *p*-value threshold of 0.05 and an absolute log_2_ fold change cutoff of at least 1 to ensure identification of genes with both statistically and biologically meaningful expression differences.

Volcano plots were generated to visualize the distribution of significant DEGs, and a Venn diagram was constructed to illustrate overlap between the DEG sets across cohorts.

### Development of RITE classifier

To ensure the classifier could be validated and used by others, TPM-normalized RNA-seq data were further normalized. For all Rodriguez TPM counts, a small epsilon of 1*e*^*−*6^ was added prior to log_2_ transformation, and values were divided by the average TPM of Rodriguez normal adjacent samples which were also added with the same small epsilon to avoid division by zero. The same procedure was applied to TPM counts from the Wang cohort^16^. Reference-based log-ratio normalization strategies have been previously used to capture biologically meaningful tumor-associated expression changes by scaling tumor expression relative to adjacent normal tissue. Prior work has demonstrated that tumor-to-normal expression ratios, computed using either patient-matched or aggregate normal references, can improve prognostic modeling and enhance interpretability in RCC and other solid tumors^24^.

We trained three supervised classifiers to distinguish among RITE, non-RITE tumor, and normal adjacent kidney tissue: multinomial logistic regression^25^, random forest^26^, and eXtreme Gradient Boosting (XGBoost)^21^. Logistic regression provided a linear baseline, while the tree-based models captured nonlinear effects and feature interactions common in tabular biomedical prediction tasks^27^.

Models were implemented in Python using scikit-learn^18^ and XGBoost (v3.0.0). Support vector machines^28^ and TabNet^29^ were also evaluated but did not outperform the selected models and were excluded from further analysis.

Logistic regression was implemented as a multinomial classifier using the limited-memory Broyden-Fletcher-Goldfarb-Shanno optimization algorithm^30^, with a maximum of 1,000 iterations specified to ensure convergence. Random forest classification was performed using an ensemble of 200 decision trees^31^, leveraging bootstrap aggregation to improve predictive accuracy and reduce overfitting through averaging across multiple tree learners.

XGBoost was trained for multiclass classification using a multiclass softmax objective that outputs per class probability estimates, with evaluation based on multiclass log loss. Hyperparameters were selected to balance model capacity and generalization. Histogram based tree construction was used for computational efficiency, tree depth was limited to 4 to constrain complexity, and boosting was performed for 300 iterations with a learning rate of 0.05^21^. Stochastic regularization was applied through row subsampling of 0.8 and per tree feature subsampling of 0.6 to reduce variance and mitigate overfitting^21^.

Model performance was evaluated across different gene feature subsets using stratified five-fold cross-validation^32^ to preserve class proportions within each fold. Multiple complementary performance metrics were computed to provide a balanced assessment of classifier behavior, particularly in the presence of class imbalance. These metrics included balanced accuracy, sensitivity, specificity, precision, F2 score, and the macro-averaged area under the receiver operating characteristic curve (AUC-ROC). Metric calculations were performed using functions from the scikit-learn library^18^.

### SHAP-based model interpretability

To improve model transparency and clinical utility, SHapley Additive exPlanation (SHAP)^22^ was used as the primary explanation method. SHAP provides both global and local interpretability through a model-agnostic, game-theoretic framework that attributes a model’s prediction to individual input features by quantifying their marginal contributions relative to a baseline. It offers consistent, additive explanations and is well-suited for both logistic regression and tree-based models.

SHAP values were computed using the TreeExplainer implementation from the SHAP library^33^. Models were trained on the optimized feature set, and class-specific SHAP values were calculated to characterize how individual genes influenced predictions for normal adjacent, non-RITE, and RITE samples.

For each model, SHAP summary plots were generated, displaying the top five features contributing to the prediction of each class, ordered by the mean magnitude of absolute SHAP values. To evaluate global importance across all classes simultaneously, absolute SHAP values were averaged within each class for every gene, producing a per-class SHAP importance matrix. A bar chart was then constructed from these values to compare the relative influence exerted by each gene across the three prediction targets.

### Screening for genes with high diagnostic utility

Using the three classifiers described above, we prioritized gene features that most strongly distinguished RITE from non-RITE tumor samples. Models were trained using both the full transcriptome and the differentially expressed gene subset identified with PyDESeq2, then interpreted using SHAP^22^ values computed with the SHAP library (version 0.50.0) in Python.

To improve robustness to any single train-test partition, we repeated stratified five-fold cross-validation across five independent random seeds, yielding 25 total splits. For each model, SHAP attributions were computed within each split, and mean absolute SHAP values were averaged across all folds and seeds to obtain a stable importance estimate per gene. Genes were then ranked by mean importance for each model, and the top 100 genes from each ranking were compared to identify consensus features shared across models.

To generate a unified prioritization of these consensus genes, we extracted each gene’s SHAP-based rank within the top 100 list for each model, where rank one indicates the most influential feature. For each gene, we computed the mean of its available ranks across models and used this mean rank to order the final consensus list, allowing genes strongly supported by multiple models to rise above those with weaker or less consistent importance.

### Determining optimal feature set

To determine the smallest gene set capable of achieving maximal classification performance, models were trained iteratively on progressively larger subsets of SHAP-ranked features. Beginning with the single highest-ranked gene and increasing sequentially to the full set of twenty-nine consensus features, each subset was evaluated using cross-validated accuracy and AUC-ROC. The optimal feature count was defined as the minimum number of genes whose inclusion produced maximal AUC-ROC for each model.

After identifying thirteen genes as the minimal set that preserved peak performance across models, this panel was compared against alternative feature subsets. These included: the top twenty-nine SHAP-ranked genes; the top twenty-nine genes ranked by absolute fold change from DESeq2; the top thirteen SHAP-ranked genes; the top thirteen DESeq2-ranked genes; and a seven-gene subset comprised of SHAP features shared by all three models.

All performance distributions were generated using twenty-fold cross-validation. Pairwise statistical comparisons were conducted using Wilcoxon rank-sum tests, with *p*-values computed in Python and annotated on boxplots using the statannotations package (version 0.7.2).

### Gene ontology and pathway enrichment analyses

Gene Ontology (GO) functional annotation and Kyoto Encyclopedia of Genes and Genomes (KEGG)^34^ pathway enrichment were performed using ShinyGO^35^ (version 0.85.1), an interactive web-based platform for gene set enrichment analysis^35^, with all expressed genes from the combined Rodriguez/Wang^16^ dataset serving as the background reference set. KEGG is a curated knowledgebase that integrates genomic, biochemical, and functional information to represent genes within well-defined metabolic and signaling pathways^34^. Bar plots of fold enrichment and false discovery rates were generated directly from ShinyGO output^35^.

To compare biological pathways shared across gene sets, enriched GO and KEGG terms were converted into gene membership lists and visualized using UpSet plots, which enable systematic identification of pathway overlaps across multiple analyses. Shared pathways identified through these intersections were further examined using KEGG pathway diagrams retrieved via ShinyGO, which accesses KEGG through API calls implemented by the pathview R package^35^. Enriched genes were mapped onto curated KEGG pathway diagrams to visualize their positions within established molecular networks, thereby contextualizing their biological relevance. Finally, chromosomal localization of all DEGs was additionally visualized to identify genomic regions exhibiting enrichment of RITE-associated transcriptional changes.

## Results

### Patient and sample characteristics

The Rodriguez cohort consisted of twenty-two patients who underwent nephrectomy with tumor thrombus resection, with a mean age of 63.6 years. Nineteen cases were ccRCC, two were papillary RCC, and one was adrenocortical carcinoma. Because RITE represents a biologic and anatomic phenotype rather than a histology-specific diagnosis, RITE samples from non-ccRCC tumors were retained in the cohort to maximize use of available data and to capture the full spectrum of venous invasion biology. All 66 non-RITE tumor samples used for downstream comparative analyses were derived from ccRCC.

Across all three datasets included in the global PCA analysis (Fig. 1), we analyzed a total of 721 bulk RNA-seq samples: 71 from the Rodriguez cohort, 174 from the Wang cohort^16^, and 476 from TCGA-KIRC. Each dataset contributed multiple tissue types relevant to RITE biology. Sample distributions for Rodriguez and Wang^16^ were:

**Figure 1.**
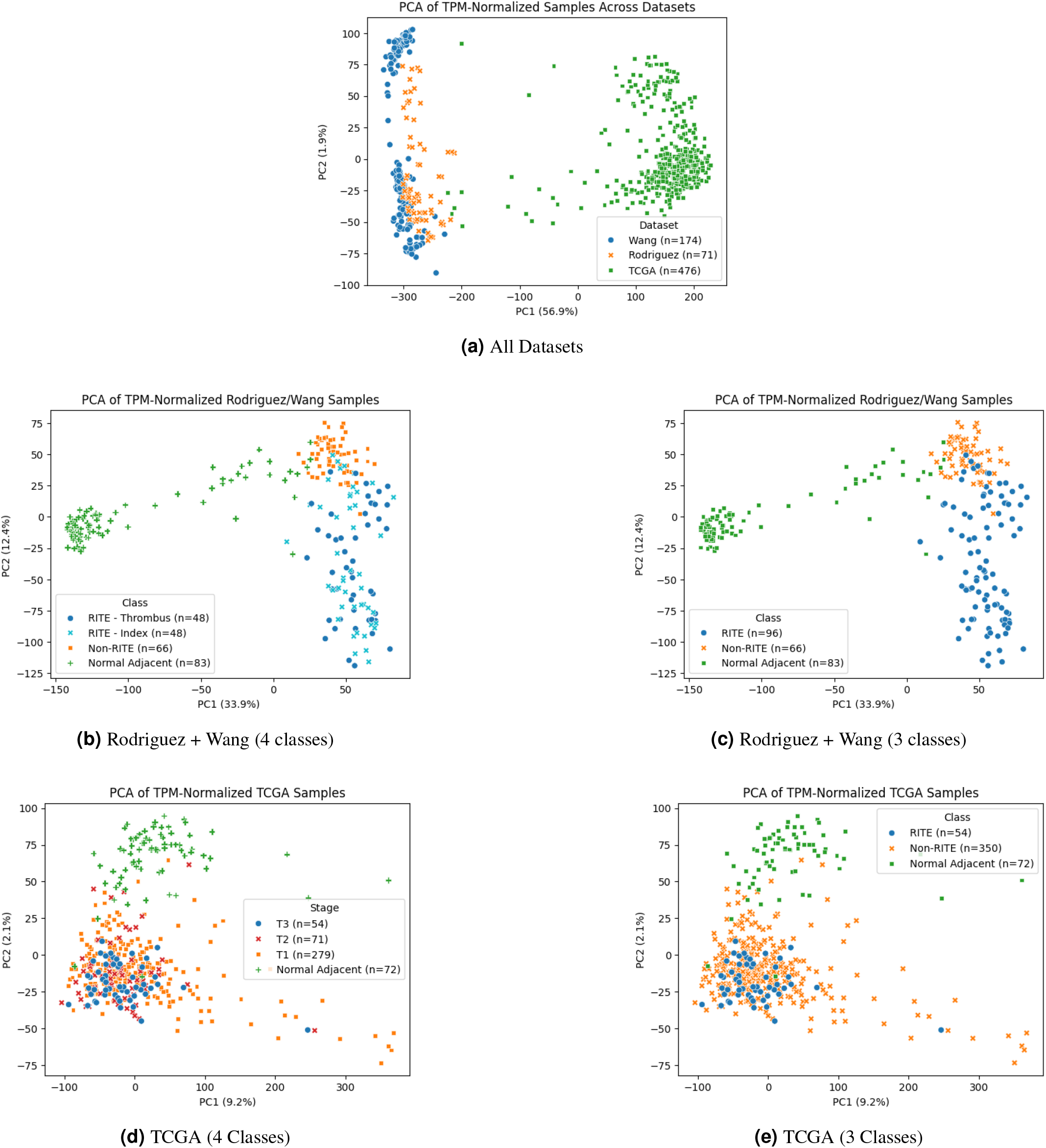
PCA of RCC transcriptomes across cohorts. **(a)** PCA of all three datasets (Rodriguez, Wang, TCGA-KIRC). **(b–c)** PCA restricted to the Rodriguez and Wang cohorts. **(d–e)** PCA of TCGA samples.

- **RITE-thrombus:** 22 Rodriguez samples, 26 Wang^16^ samples
- **RITE-index (non-vascular portion of tumor from patients with thrombus):** 22 Rodriguez, 26 Wang^16^ samples
- **RITE-normal adjacent tissue:** 22 Rodriguez, 61 Wang^16^ samples
- **Non-RITE tumors:** 5 Rodriguez, 61 Wang^16^ samples

TCGA-KIRC contained only tumor and normal adjacent samples. When aligned to our classification framework, TCGA contributed 54 RITE tumors, 350 non-RITE tumors, and 72 normal adjacent kidney tissues.

These distributions provided adequate representation across RITE and non-RITE states, enabling robust differential expression, pathway enrichment, and cross-cohort validation analyses.

### Transcriptomic profiling of RITE

TPM-normalized RNA-seq data spanning 45,408 genes and 721 samples from the Rodriguez, Wang^16^, and TCGA-KIRC cohorts were used for dimensionality-reduction analyses. Within each dataset, genes with an average raw count below one were removed to reduce low-abundance noise, resulting in 17,658 retained genes for Rodriguez, 17,031 for Wang^16^, and 13,003 for TCGA. Cross-cohort PCA was performed using the 12,672 genes shared among all three datasets after log-transformation using log_2_(TPM + 1*e*^−6^).

Cross-cohort PCA revealed that TCGA samples formed a distribution distinct from the Rodriguez and Wang^16^ datasets (Fig. 1a), indicating substantial batch or cohort effects that precluded integrated downstream analysis. Therefore, subsequent PCA was conducted separately for the two-cohort Rodriguez/Wang^16^ and TCGA datasets.

For the Rodriguez and Wang^16^ cohorts, PCA performed using the intersection of 16,623 shared genes demonstrated a consistent structure across both four-class—RITE-thrombus, RITE-index, non-RITE, normal adjacent—and three-class—RITE, non-RITE, normal adjacent—analyses. Normal adjacent tissue formed a well-defined cluster distinct from tumor samples, while RITE-thrombus and RITE-index tumors overlapped extensively, reflecting strong transcriptional similarity between the primary tumor and its associated tumor extension. Non-RITE tumors occupied a partially separate region of the PCA space, indicating biologically meaningful divergence, although some overlap with RITE tumors persisted (Fig. 1b–c).

In contrast, PCA of the TCGA dataset revealed a fundamentally different structure as expected. While normal adjacent tissue again formed a discrete cluster, T1, T2, and T3 tumors showed extensive overlap with no discernible stage-dependent separation. When T3BC tumors were reclassified as RITE and T1/T2 as non-RITE, the groups remained transcriptionally indistinguishable, suggesting that TCGA-KIRC does not capture the molecular features associated with bona fide RITE (Fig. 1d-e). These findings support restricting differential expression and downstream modeling to the Rodriguez and Wang^16^ datasets, which together demonstrated internally consistent RITE-related biology.

Differential expression analysis was performed using raw RNA-seq counts from all available samples. Because PyDESeq2 applies its own normalization procedures, TPM-normalized values were excluded from this step. To reduce noise from low-abundance transcripts, genes with an average raw count below one were removed independently for each dataset. After filtering, the combined Rodriguez and Wang^16^ cohorts retained 26,474 genes across 245 samples, whereas TCGA retained 11,165 genes across 476 samples. PCA of the merged Rodriguez/Wang^16^ raw counts showed no cohort-driven separation, supporting the use of a combined matrix without explicit batch correction.

Within each dataset, differential expression testing compared RITE tumors with non-RITE tumors. For Rodriguez/Wang^16^, this included 96 RITE and 66 non-RITE tumors. In TCGA, tumor stage served as a surrogate for thrombus biology, with T3BC classified as RITE (n = 54) and T1/T2 as non-RITE (n = 350). DESeq2 was applied using an adjusted *p*-value threshold of 0.05 and an absolute log_2_ fold change *≥* 1. The combined Rodriguez and Wang^16^ datasets yielded 6,317 significant DEGs with 3,705 upregulated and 2,612 downregulated, representing a broad and biologically coherent transcriptional shift associated with RITE (Fig. 2a). The volcano plot revealed a wide distribution of effect sizes, consistent with large-scale transcriptomic reprogramming in tumors with venous invasion.

**Figure 2.**
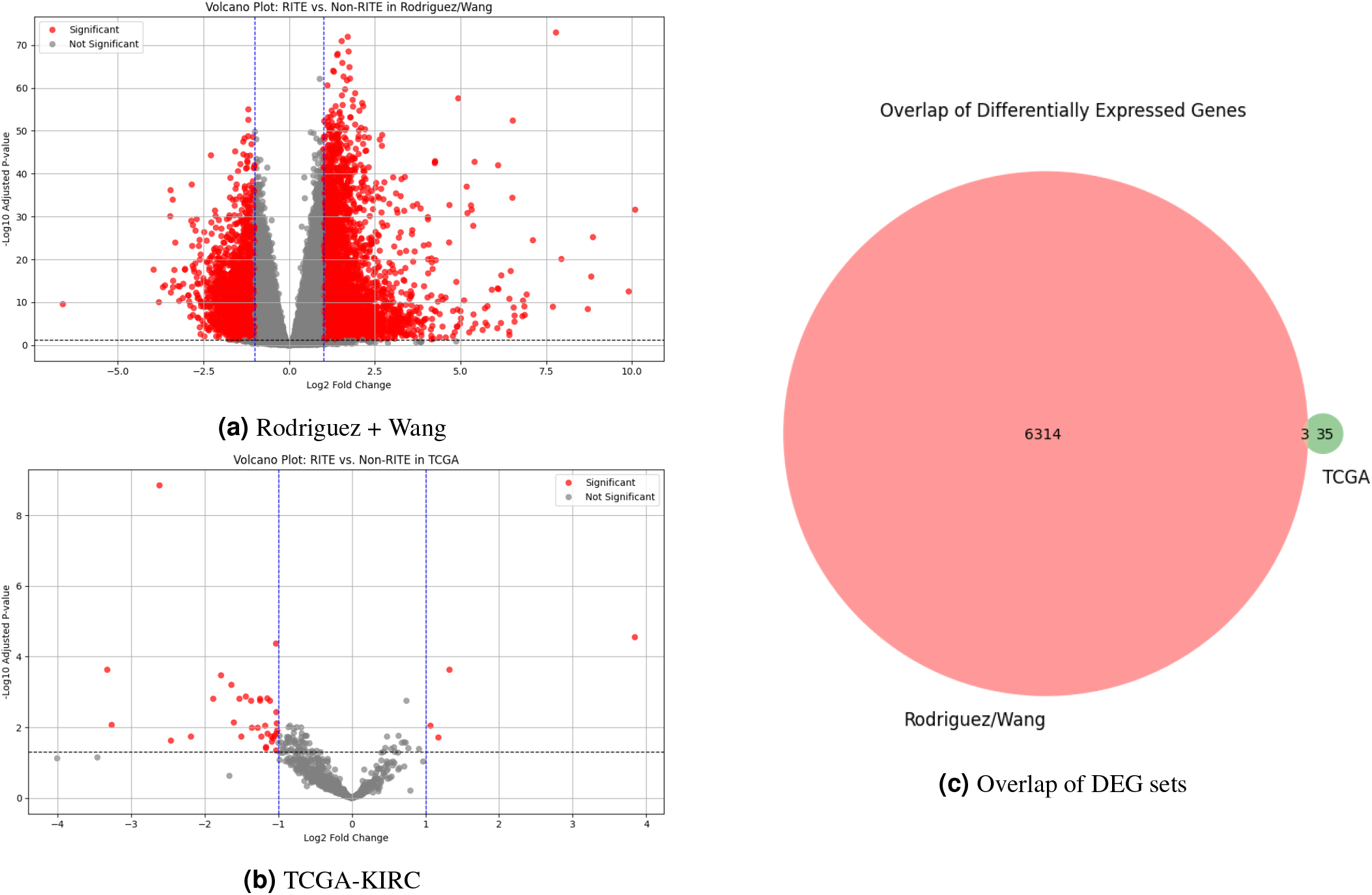
Differential gene expression in RITE versus non-RITE tumors. (a) Volcano plot of DEGs in the combined Rodriguez and Wang datasets. (b) Volcano plot of DEGs in TCGA-KIRC using T3BC as a surrogate for RITE. (c) Venn diagram comparing DEG overlap between the two datasets; only three genes were shared.

By contrast, TCGA-KIRC produced only 38 significant DEGs under the same thresholds with 4 upregulated and 34 downregulated, with a markedly restricted distribution of fold changes (Fig. 2b). This limited signal reflects both the use of pathologic stage as an imperfect surrogate for RITE and the broader absence of RITE-specific molecular information in TCGA, as observed in PCA.

Comparison of DEG lists revealed only three genes in common—*EGR1, FXYD2*, and *MT-ATP8* (Fig. 2c). None of these were regulated concordantly across datasets, further indicating that TCGA does not capture the transcriptional landscape in sufficient depth to reveal the detailed molecular program underlying true venous tumor thrombus.

### RITE prediction using an intravascular extension-associated gene classifier

To identify a parsimonious gene signature capable of distinguishing RITE from non-RITE tumors, we evaluated three supervised learning algorithms chosen for their strong performance on high-dimensional tabular data: logistic regression, random forest, and XGBoost.

All models were initially trained on the 26,474 genes with mean raw count greater than one in the combined Rodriguez/Wang^16^ dataset. Baseline experiments using this full feature set established an upper bound on performance. Restricting training to the 6,317 DEGs identified by DESeq2 improved or preserved classification metrics relative to the full gene set (Table 1), supporting the use of DEG-filtered features for subsequent refinement.

**Table 1.** Classifier performance across gene feature subsets. Balanced accuracy (Bal Acc), sensitivity (Sens), specificity (Spec), precision (Prec), F_2_, and macro AUC-ROC (AUC) for logistic regression, random forest, and XGBoost trained on four feature sets: all expressed genes, all DEGs, the 29-gene SHAP consensus panel, and the 13-gene minimal SHAP panel. Bold values indicate the best-performing feature set for each metric within each model, highlighting the benefit of SHAP-guided dimensionality reduction.

| Feature set | Logistic Regression |  |  |  |  |  | Random Forest |  |  |  |  |  | XGBoost |  |  |  |  |  |
| --- | --- | --- | --- | --- | --- | --- | --- | --- | --- | --- | --- | --- | --- | --- | --- | --- | --- | --- |
|  | Bal Acc | Sens | Spec | Prec | F2 | AUC | Bal Acc | Sens | Spec | Prec | F2 | AUC | Bal Acc | Sens | Spec | Prec | F2 | AUC |
| All 26,474 genes | 0.964 | 0.964 | 0.983 | 0.972 | 0.964 | 0.997 | 0.960 | 0.960 | 0.981 | 0.967 | 0.960 | 0.999 | 0.977 | 0.977 | 0.989 | 0.982 | 0.978 | 0.998 |
| 6,317 DEGs | 0.976 | 0.976 | 0.989 | 0.984 | 0.977 | 0.999 | 0.964 | 0.964 | 0.983 | 0.972 | 0.964 | 0.999 | 0.974 | 0.974 | 0.987 | 0.978 | 0.974 | 0.998 |
| 29 consensus genes | **0.986** | **0.986** | **0.994** | **0.990** | **0.987** | **1.000** | 0.976 | 0.976 | 0.989 | 0.984 | 0.977 | **1.000** | **0.988** | **0.988** | **0.994** | **0.988** | **0.988** | **1.000** |
| 13 selected genes | 0.978 | 0.978 | 0.990 | 0.981 | 0.978 | **1.000** | **0.981** | **0.981** | **0.991** | **0.986** | **0.982** | **1.000** | 0.983 | 0.983 | 0.992 | 0.984 | 0.983 | **1.000** |

To further prioritize informative features, we computed SHAP values for each model and averaged them across five-fold cross-validation repeated over five random seeds. Genes were ranked by mean absolute SHAP value for each algorithm, and the top 100 features per model were compared. SHAP bar plots illustrate the highest-importance genes for logistic regression, random forest, and XGBoost, respectively (Fig. 3a–c). The overlap analysis of these top 100 sets yielded 29 consensus genes present in at least two models, including 7 genes shared by all three classifiers (Fig. 3d). These consensus features and their cross-model ranks are summarized in Table 2. Retraining the classifiers using only this 29-gene panel improved accuracy and AUC-ROC compared with models trained on all expressed genes or all DEGs (Table 1).

**Table 2.** Consensus SHAP feature rankings across classifiers. The 29-gene consensus set was defined as genes appearing among the top 100 SHAP-ranked features in at least two of three models (logistic regression, random forest, XGBoost). For each gene, the number of models in which it appeared (Models used), its best single-model rank, mean rank across models, and mean absolute SHAP value are shown.

| Feature | Models used | Best rank | Mean rank | Mean SHAP |
| --- | --- | --- | --- | --- |
| FOXI1 | 3 | 2 | 3.0 | 0.0683 |
| PRR35 | 2 | 1 | 4.5 | 0.0256 |
| GP2 | 3 | 2 | 6.7 | 0.0298 |
| CLDN8 | 3 | 1 | 13.3 | 0.0636 |
| KNG1 | 2 | 9 | 16.5 | 0.0164 |
| HEPACAM2 | 3 | 11 | 16.7 | 0.0259 |
| SALL3 | 2 | 5 | 17.0 | 0.0291 |
| FXYD4 | 2 | 14 | 17.0 | 0.0175 |
| BSND | 3 | 1 | 17.3 | 0.2458 |
| ATP6V1G3 | 3 | 6 | 17.3 | 0.0233 |
| SNORD15B | 2 | 2 | 19.0 | 0.1479 |
| TFAP2B | 3 | 16 | 20.7 | 0.0224 |
| ATP6V0A4 | 2 | 19 | 22.0 | 0.0137 |
| TMEM213 | 2 | 20 | 25.5 | 0.0112 |
| AC092640.1 | 2 | 6 | 29.0 | 0.0500 |
| AQP2 | 2 | 4 | 32.5 | 0.0090 |
| RHBG | 2 | 11 | 32.5 | 0.0081 |
| RSC1A1 | 2 | 9 | 36.5 | 0.0370 |
| UMOD | 2 | 21 | 38.5 | 0.0157 |
| KCNJ1 | 2 | 3 | 39.5 | 0.0082 |
| ATP12A | 2 | 14 | 42.5 | 0.0168 |
| VWA5B1 | 2 | 21 | 43.5 | 0.0060 |
| PIK3C2G | 2 | 15 | 47.5 | 0.0165 |
| AC006486.1 | 2 | 7 | 51.5 | 0.0203 |
| ZIC1 | 2 | 10 | 51.5 | 0.0190 |
| FABP6 | 2 | 30 | 56.5 | 0.0042 |
| RETN | 2 | 46 | 70.0 | 0.0140 |
| AC020661.4 | 2 | 64 | 71.0 | 0.0039 |
| AC009137.2 | 2 | 74 | 80.5 | 0.0135 |

**Figure 3.**
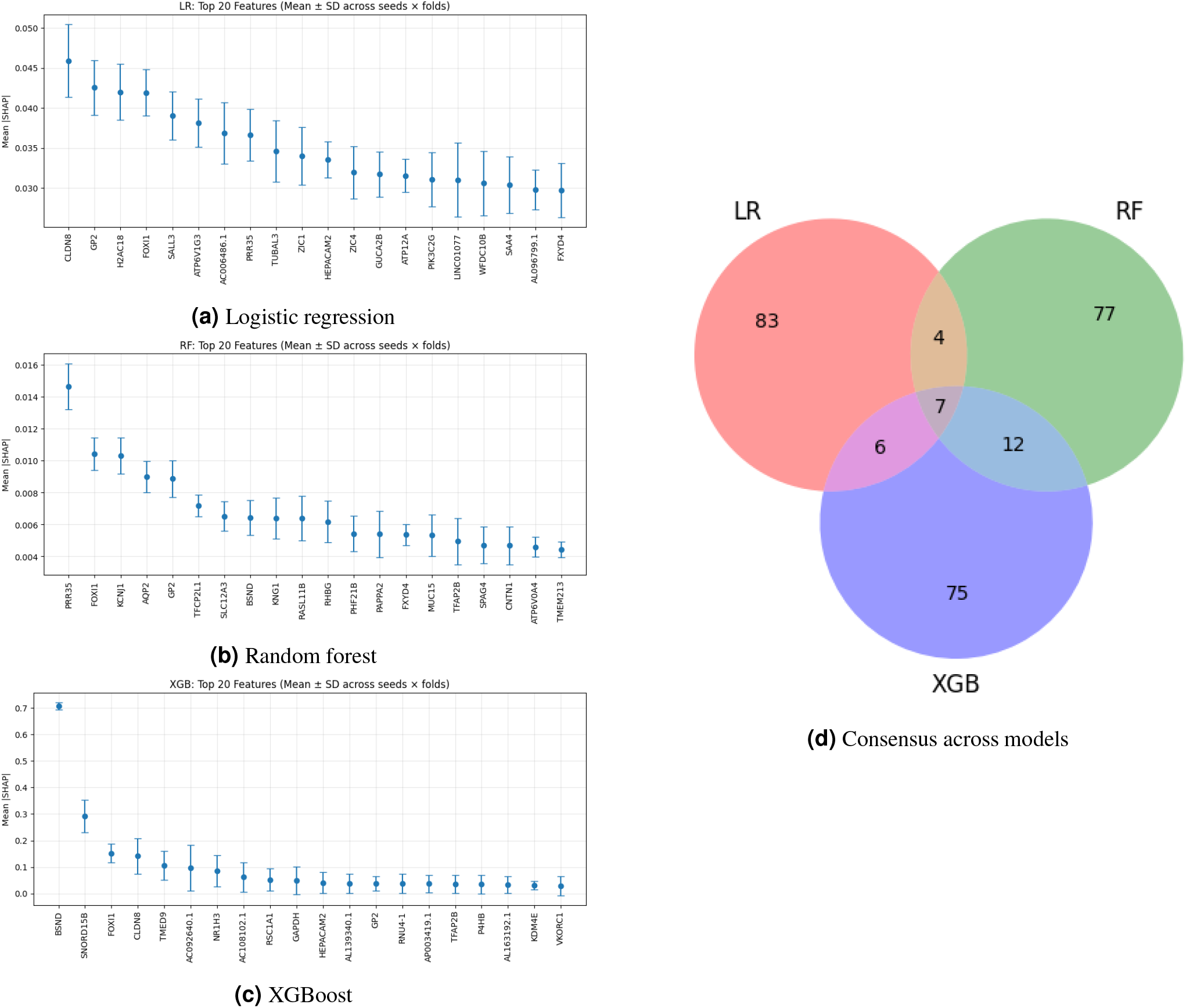
SHAP-based feature importance and consensus genes across classifiers. (a–c) Mean SHAP values for the top-ranked genes in logistic regression, random forest, and XGBoost models trained to classify RITE versus non-RITE tumors, illustrating partially overlapping but model-specific importance profiles. (d) Venn diagram showing the intersection of the top 100 SHAP-ranked features from each model, yielding a 29-gene consensus set, including 7 genes shared by all three classifiers.

To determine the minimal number of genes required to preserve maximal performance, we conducted a greedy ranked forward feature inclusion procedure^36,37^ guided by global SHAP importance^38,39^. For each model, features were added sequentially from the highest-to lowest-ranked SHAP gene within the 29-gene consensus set, and performance was evaluated at each step by cross-validated accuracy and AUC-ROC (Fig. 4a–c). Performance curves plateaued once the top-ranked genes were included: logistic regression and random forest reached peak accuracy and AUC-ROC at 13 genes, whereas XGBoost achieved its maximal AUC-ROC with 11 genes but showed no further improvement when extended to 13. We therefore adopted a uniform 13-gene minimal panel for all classifiers.

**Figure 4.**
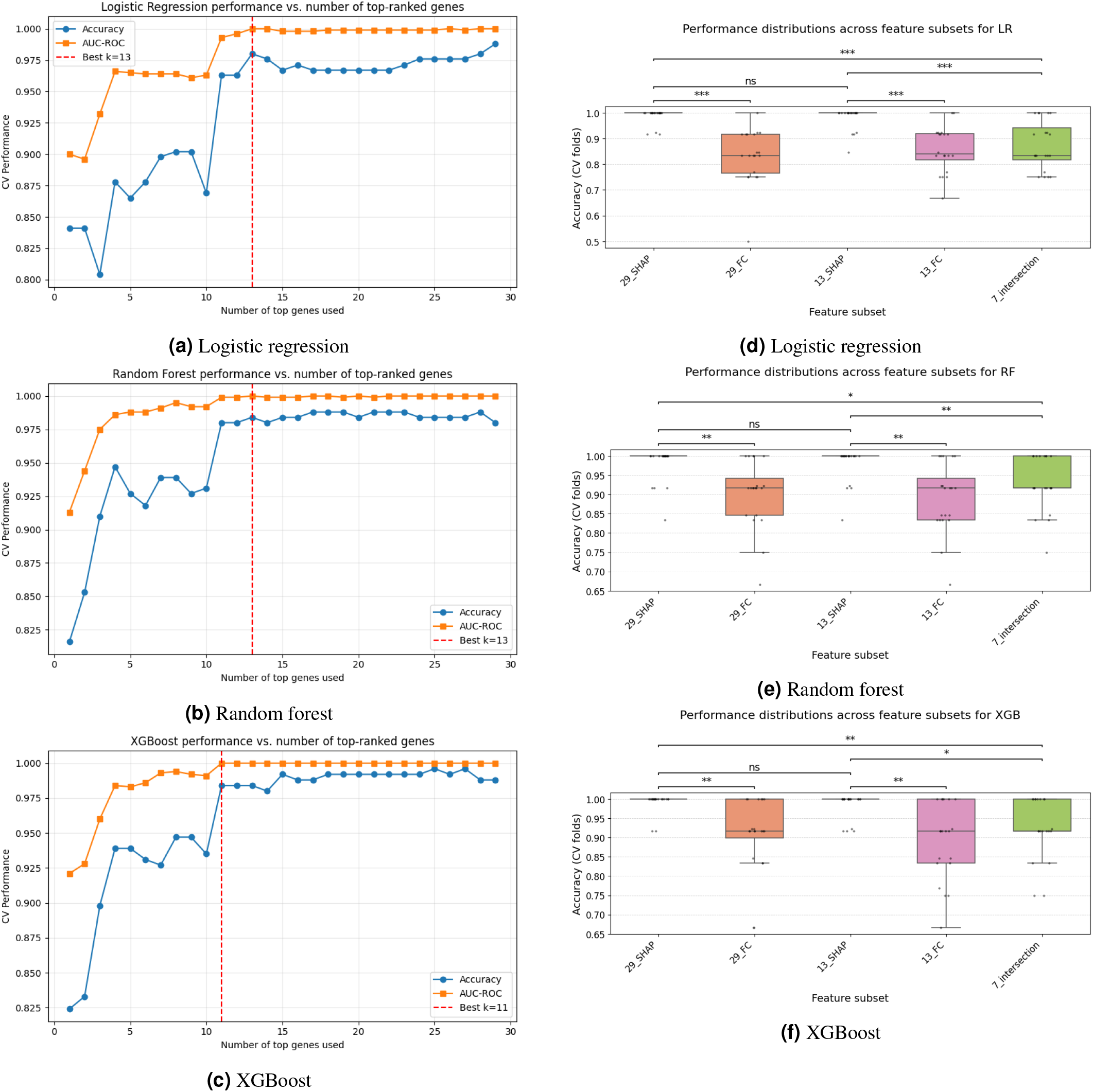
Greedy SHAP-based feature selection and comparison of gene subsets. (a–c) Greedy inclusion curves showing cross-validated classification performance as SHAP-ranked genes from the 29-gene consensus panel are added sequentially to logistic regression, random forest, and XGBoost models. Performance plateaus once the highest-ranked genes are included, with maximal accuracy and AUC-ROC achieved using 11–13 features. (d–f) Boxplots comparing cross-validated accuracy for multiple feature subsets, including all expressed genes, all DEGs, the 29-gene consensus set, the 13-gene minimal SHAP panel, DESeq2 rank-based subsets, and the 7-gene intersection shared by all models, demonstrating consistently superior performance of SHAP-guided gene selections.

We next compared this 13-gene panel against several alternative feature subsets: the full set of 29 consensus genes, the top 29 and top 13 genes ranked solely by absolute fold change from DESeq2, and the 7-gene intersection shared by all three models. Using twentyfold cross-validation, the 13- and 29-gene SHAP-based subsets consistently outperformed their DESeq2-only counterparts across classifiers (Fig. 4d–f). There was no significant gain in performance when increasing from 13 to 29 SHAP-ranked genes, whereas the 7-gene intersection panel showed a noticeable drop in accuracy and AUC-ROC, indicating that the broader 13-gene set was necessary to maintain optimal predictive power.

Model performance across feature sets is summarized in Table 1. For logistic regression and XGBoost, the 29-gene consensus panel produced the highest overall metrics, with balanced accuracy up to 0.988 and macro AUC-ROC of 1.000, while the 13-gene panel achieved comparable AUC-ROC with only a modest change in other metrics. Random forest achieved its best balanced accuracy, sensitivity, specificity, precision, F_2_, and macro AUC-ROC when trained on the 13-gene panel. Across all three algorithms, models trained on either all expressed genes or all DEGs underperformed compared with SHAP-guided feature subsets, underscoring the benefit of interpretable, data-driven feature selection.

Finally, we used SHAP to interpret how individual genes influenced class-specific predictions in the best-performing tree-based models. Random forest and XGBoost trained on the 13-gene panel were subjected to class-wise SHAP analysis for normal adjacent, non-RITE, and RITE samples. SHAP summary plots highlight the top features driving each class prediction (Fig. 5a–c,e–g), and global bar plots show average absolute SHAP values across classes (Fig. 5d,h). Across both models, *FOXI1* emerged as a dominant feature for normal adjacent classification, whereas *SNORD15B* was particularly influential in separating non-RITE and RITE tumors. Additional genes such as *GP2, CLDN8, ATP6V0A4, PRR35*, and *BSND* contributed more modest but consistent class-dependent effects. Together, these results define a unique and compact, biologically interpretable gene signature that robustly predicts RITE status from tumor transcriptomes.

**Figure 5.**
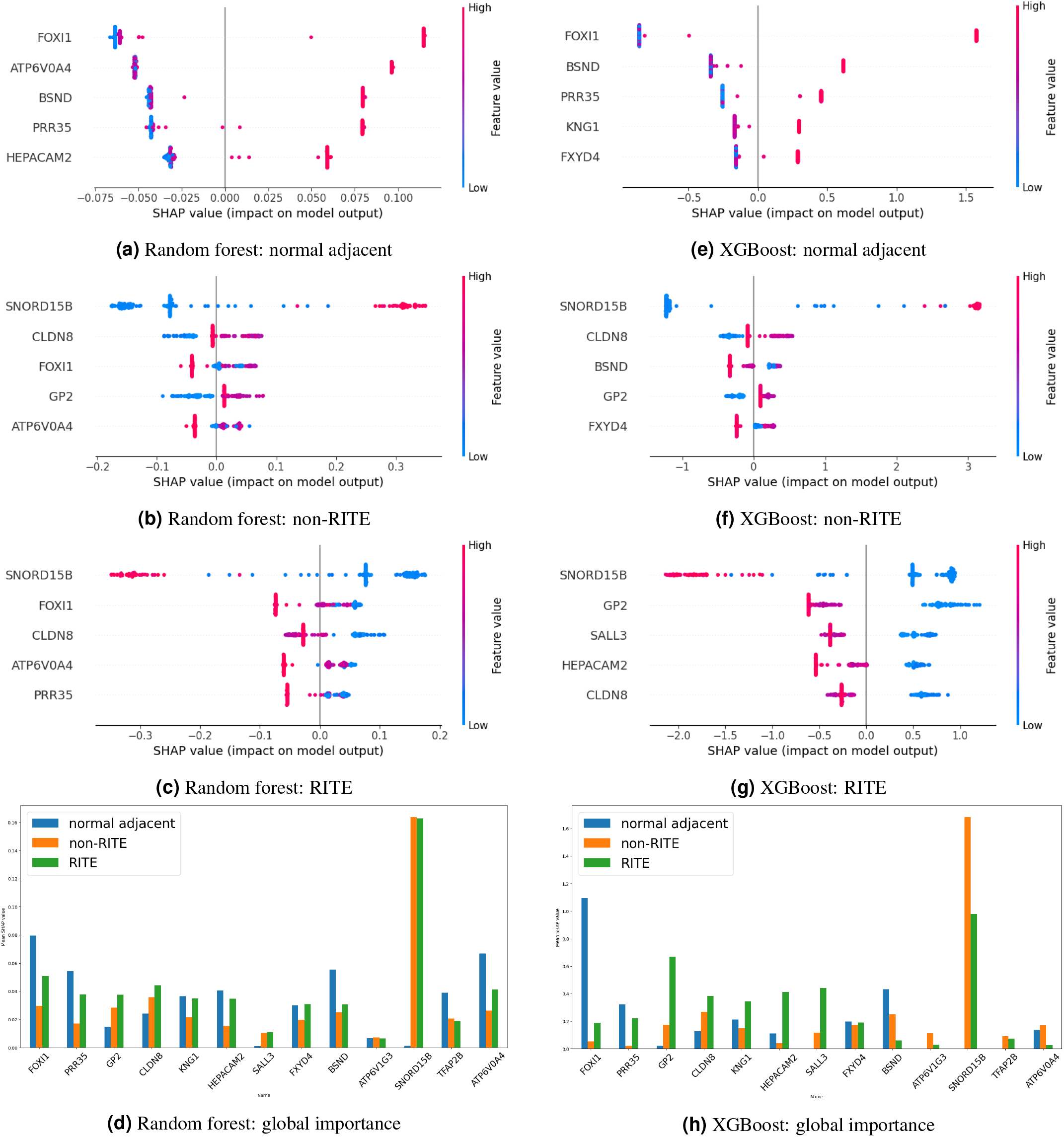
Class-specific and global SHAP interpretations for tree-based RITE classifiers. (a–c, e–g) Class-wise SHAP summary plots for random forest and XGBoost models trained on the 13-gene panel, showing the top features driving predictions of normal adjacent, non-RITE, and RITE classes. Each point represents a sample, with color indicating gene expression level and position on the *x*-axis denoting the SHAP contribution to the predicted class. (d, h) Global SHAP bar plots summarizing average absolute SHAP values across all classes, highlighting *FOXI1, SNORD15B*, and additional genes such as *GP2, CLDN8*, and *BSND* as key contributors to model predictions.

### Pathway and gene ontology enrichment

Pathway and gene ontology enrichment analyses were performed using ShinyGO (version 0.85.1)^35^, with the 26,474 expressed genes from the Rodriguez/Wang^16^ dataset serving as the background. Four gene sets were evaluated: all 6,317 DEGs, the 3,705 upregulated DEGs, the 29 SHAP-derived consensus genes, and the optimized 13-gene panel. For each set, ShinyGO-generated bar plots of fold enrichment and false discovery rate (FDR) were examined (Fig. 6a–d)^35^.

**Figure 6.**
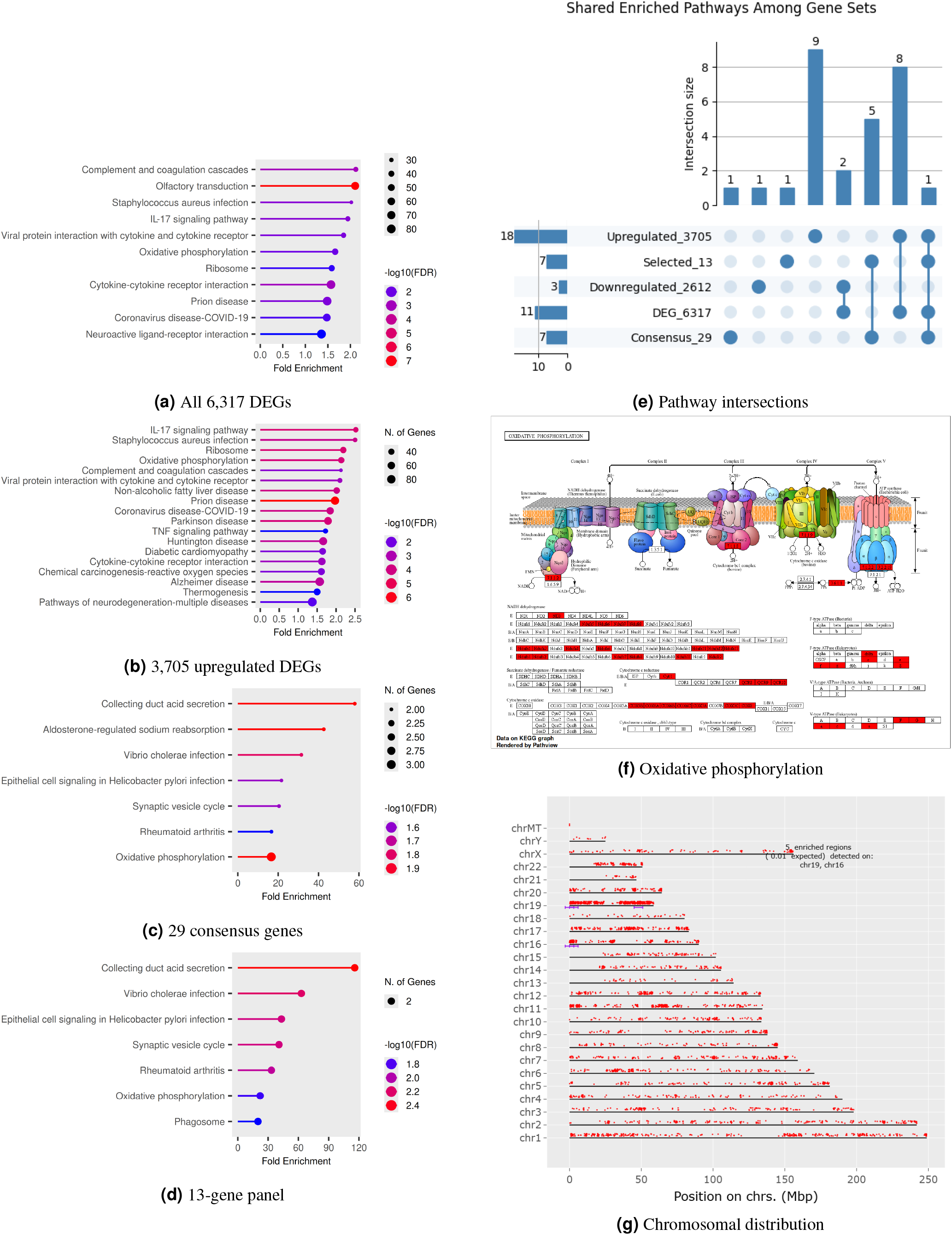
Pathway and gene ontology enrichment across RITE-associated gene sets. (a–d) ShinyGO pathway and gene ontology enrichment for all 6,317 DEGs, the 3,705 upregulated DEGs, the 29 SHAP-derived consensus genes, and the optimized 13-gene panel, using the 26,474 expressed genes in the Rodriguez/Wang dataset as background. (e) UpSet plot summarizing intersections among enriched pathways across the four gene sets. (f) KEGG oxidative phosphorylation diagram highlighting pathway genes represented within the enriched sets. (g) Chromosomal mapping of all DEGs, showing the distribution of differentially expressed genes across the genome.

Enrichment of the full DEG list revealed strong overrepresentation of pathways related to complement and coagulation cascades, cytokine signaling, and metabolic processes (Fig. 6a). When restricted to the 3,705 upregulated DEGs, the enrichment profile broadened, with IL-17 signaling exhibiting the highest fold enrichment and additional immune and metabolic pathways emerging more prominently after directional filtering (Fig. 6b).

The 29-gene SHAP consensus set produced a more focused enrichment pattern, in which collecting duct acid secretion and other epithelial transport-related pathways featured prominently, along with pathways linked to mitochondrial function (Fig. 6c). The optimized 13-gene panel largely preserved these signals, with collecting duct acid secretion again ranking as the top pathway, indicating that the reduced feature set retained key mechanistic information associated with RITE biology (Fig. 6d).

To compare biological themes shared among gene sets, enriched pathways were converted into gene set memberships and visualized using an UpSet plot (Fig. 6e). Oxidative phosphorylation emerged as a consistently shared pathway across multiple gene sets, and inspection of the KEGG oxidative phosphorylation diagram confirmed that several components of the respiratory chain were represented among the enriched genes (Fig. 6f), reinforcing the centrality of mitochondrial metabolism in the RITE-associated transcriptomic program.

Finally, chromosomal mapping of all 6,317 DEGs revealed non-uniform genomic distribution, with notable concentrations on chromosomes 16 and 19 (Fig. 6g), suggesting that RITE-associated transcriptional changes may be influenced by cluster-level genomic organization not previously associated with RCC.

## Discussion

We describe a large molecular analysis of the gene expression profiles between RITE versus similar sized RCC without vascular involvement. The analysis provides substantial insight into the gene programming derangements that occur in this unique phenotype.

PCA provided a robust framework for assessing dataset compatibility and identifying transcriptomic structure relevant to venous tumor thrombus. The Rodriguez and Wang^16^ datasets demonstrated strong concordance, clustering together and enabling joint downstream analysis (Fig. 1b–c). This agreement likely reflects similarities in sequencing platforms, sample handling, and data processing pipelines across the two studies. Within this combined space, normal adjacent tissues separated cleanly from tumor samples, validating sample integrity and preprocessing choices. The close overlap between RITE-index and RITE-thrombus specimens further supports their shared cellular origin and transcriptional program, consistent with the anatomical relationship between thrombus and its parent tumor. Although Shi et al. reported differences between index and thrombus tissues^40^, the near-complete overlap observed here indicates that such differences are small relative to the global RITE/non-RITE distinction and were not pursued further.

In contrast, TCGA-KIRC did not reproduce these transcriptional signatures. Despite its large sample size, TCGA tumors showed no meaningful separation by stage or by RITE-surrogate labeling (Fig. 1d–e). These discrepancies likely reflect substantial technological and temporal differences: TCGA was sequenced more than a decade earlier using older, lower-depth platforms, resulting in limited dynamic range and reduced sensitivity for low-abundance transcripts. Because the dataset predates dedicated RITE annotation and lacks true tumor thrombus samples, stage-based surrogates (e.g., T3BC vs. T1/T2) may not accurately capture biological processes specific to venous extension. Together, these observations justified restricting differential expression and pathway analyses to the Rodriguez and Wang^16^ datasets, where internally consistent transcriptomic patterns aligned with known disease biology.

Differential expression analysis revealed a striking contrast between datasets. The combined Rodriguez/Wang^16^ cohort yielded 6,317 DEGs separating RITE from non-RITE tumors, consistent with the clear PCA distinction and the inclusion of true thrombus and matched tumor tissues (Fig. 2a). In comparison, TCGA produced only 38 DEGs under equivalent thresholds (Fig. 2b), reflecting both technological limitations of early RNA-seq analysis and the low abundance of RITE tumors. The minimal three-gene overlap—*EGR1, FXYD2, MT-ATP8*—combined with discordant fold-change directions (Fig. 2c), underscores that TCGA cannot serve as a surrogate dataset for studying venous tumor extension. By contrast, the Rodriguez/Wang^16^ datasets capture a biologically relevant transcriptional signature that distinguishes RITE from non-RITE disease.

Integrating SHAP-based feature attribution across three machine learning models enabled identification of a 29-gene consensus set that consistently contributed to RITE classification (Fig. 3). The strong agreement among logistic regression, random forest, and XGBoost indicates that these genes reflect robust biological signals rather than model-specific artifacts. Reducing from 26,474 expressed genes to a concise consensus set improved predictive accuracy while substantially simplifying model complexity (Table 1). The approach highlights the utility of interpretable machine learning in high-dimensional transcriptomic settings, where convergence across independent models can guide the identification of biologically meaningful and reproducible features.

*FOXI1* emerged as the most influential classifier gene across models, consistent with its known role in renal epithelial differentiation and ion transport. *FOXI1* encodes a forkhead transcription factor that regulates genes essential for acid-base homeostasis, including V-ATPase subunits, pendrin, and carbonic anhydrase II^41,42^. These pathways are characteristic of *α*-intercalated cells of the distal nephron^43,44^. These cells belong to a conserved family of forkhead-related epithelial cells, which are defined by high mitochondrial content and specialized ion transport functions across multiple tissues, including the kidney and endolymphatic sac^41,45^. Its downregulation in RITE tumors suggests loss of distal epithelial identity, aligning with the expectation that ccRCC arise from more proximal tubular segments. Conversely, chromophobe RCC and oncocytoma— tumors of distal epithelial origin—are characterized by high *FOXI1* expression^46,47^, supporting the biological coherence of our findings.

The consensus panel also contained multiple epithelial transporters, ion channels, and metabolic regulators, indicating that tubular identity, electrolyte handling, and cellular energetics are central determinants of RITE-associated biology. Notably, the inclusion of *SNORD15B* among top classifier features suggests a role for altered RNA-processing or ribosomal regulation in RITE progression, consistent with emerging evidence across multiple cancer types linking small nucleolar RNAs to tumor aggressiveness and metabolic adaptation^48,49^. The aggregation method used for consensus ranking balanced model-specific sensitivity with cross-model robustness, producing a stable, high-confidence set suitable for downstream mechanistic work.

Feature selection experiments demonstrated that a compact 13-gene subset was sufficient to preserve maximal classification performance across all three classifiers (Fig. 4). The absence of performance loss when reducing from 29 to 13 genes indicates that the core discriminatory biology is concentrated within a small number of transcripts. SHAP-ranked features consistently outperformed DESeq2 fold-change-based subsets, likely because SHAP incorporates nonlinear effects and interactions that univariate differential expression analysis cannot capture. This compact gene set provides a practical foundation for translational applications, including targeted assays, biomarker development, and predictive modeling aimed at identifying RITE risk in clinical settings.

The class-specific SHAP analyses further clarified the biological roles of key genes. *FOXI1* consistently drove predictions of normal adjacent tissue, reflecting intact epithelial identity (Fig. 5a,e). *SNORD15B*, by contrast, distinguished non-RITE from RITE tumors through stepwise decreases in expression (Fig. 5b–d,f–h), suggesting progressive alterations in ribosomal or RNA-processing pathways along the trajectory from normal epithelium to RITE progression. Importantly, the most informative classifier genes were not those with the largest fold changes, but those with monotonic, class-separating behavior across normal adjacent, non-RITE, and RITE categories. This observation underscores a key distinction between univariate DEG analysis and multivariate predictive modeling.

The gene ontology analyses revealed that the full DEG set was enriched for broad tumor-associated pathways—including complement and coagulation cascades, cytokine signaling, and metabolic reprogramming^50^—mirroring prior studies of RCC progression (Fig. 6a)^51,52^. Directional filtering of upregulated DEGs amplified immune and inflammatory signatures (Fig. 6b) supported in previous studies^53^. In contrast, the consensus and minimal SHAP gene sets produced a focused enrichment profile dominated by epithelial ion transport and mitochondrial pathways (Fig. 6c–d). Oxidative phosphorylation emerged as a consistently enriched pathway across all gene sets (Fig. 6e–f), highlighting its central role in RITE-associated biology. This finding is concordant with prior literature showing mitochondrial alterations and metabolic reprogramming in RCC^54–56^, including the work of Shi et al. identifying oxidative phosphorylation upregulation in both tumorigenesis and thrombus invasion^57^.

Chromosomal mapping further suggested genomic structure underlying differential expression, with enrichment on chromosomes 16 and 19 (Fig. 6g). Prior mutation studies, including those by Wang et al., have similarly reported structural alterations associated with venous invasion^16^, suggesting potential interplay between genomic and transcriptomic determinants of thrombus formation^58^.

Collectively, the PCA, DEG, SHAP, and gene ontology analyses converge on a coherent biological narrative: RITE tumors exhibit distinct transcriptional programs characterized by loss of distal epithelial identity, reconfiguration of ion transport machinery, and consistent perturbation of mitochondrial metabolism. Taken together, these findings support a model in which *FOXI1* downregulation, altered RNA-processing programs, and oxidative phosphorylation remodeling collectively contribute to venous tumor extension. The resulting 13-gene panel represents a compact, biologically interpretable signature capable of robustly distinguishing RITE, non-RITE, and normal adjacent tissue. This work provides a foundation for mechanistic investigation into venous tumor extension and offers a practical, data-driven framework for future diagnostic and prognostic tool development in RCC. Further mechanistic investigation of these molecular pathways may broaden the applicability of our RITE classifier and help inform clinical management strategies.

We recognize several limitations of this study. Most notably, the analyses are retrospective and lack direct clinical validation. In addition, TCGA was unable to contribute to the final classifier, as the limited dynamic range of the RNA-seq technology used in TCGA constrains its ability to capture low-abundance transcripts relevant to RITE biology. Because of the difficulty in collecting high-quality matched normal, tumor, and thrombus specimens, RNA-seq was performed on 71 tissue samples from 22 patients in the Rodriguez cohort. This sample size was sufficient to detect transcriptional differences at the sequencing level but did not allow for complete one-to-one matching across tissue types. Furthermore, the cohort included a small number of non-ccRCC cases as the primary focus was RITE versus non-RITE, rather than restricting to a single histologic subtype, though all but four samples were ccRCC as this is the most common RCC subtype. Given the limited sample size, the prevalence of some potentially informative genes did not differ significantly among normal, tumor, and thrombus tissues. Finally, this study was exploratory in nature and aimed to provide a global view of the molecular features associated with RITE. Further studies are required to define the biological functions of these genes and to determine whether the observed transcriptional changes directly contribute to intravascular tumor extension.

Despite these limitations, our results highlight the pivotal roles of *FOXI1, SNORD15B*, and oxidative phosphorylation in RITE development in RCC patients. Given the urgent unmet need for improved risk stratification and early identification of venous tumor extension, these findings support the future development of a biologically informed prognostic panel. Subsequent studies will focus on integrating clinicopathologic and survival data and validating these signatures using immunohistochemical staining of human RCC tissues. External validation in independent cohorts will be required to establish clinical applicability.

## Conclusion

This study integrates transcriptomic profiling, machine learning feature attribution, and pathway analysis to define the molecular landscape underlying venous tumor extension in renal cell carcinoma. PCA demonstrated that only the Rodriguez and Wang^16^ datasets contained coherent RITE-associated structure, enabling reliable differential expression and model-based feature discovery. Using these datasets, we identified 6,317 DEGs distinguishing RITE from non-RITE tumors and derived a reproducible 29-gene consensus set through SHAP-guided attribution across three independent classifiers. Further refinement yielded a compact 13-gene panel capable of matching or exceeding the performance of substantially larger feature sets, underscoring that the core biology of venous extension resides within a small number of highly informative transcripts, including *FOXI1* and *SNORD15B*.

The biological themes identified—loss of distal epithelial identity, perturbation of ion transport machinery, and consistent enrichment of mitochondrial pathways—highlight mechanistic programs that may drive the aggressive behavior of RITE tumors. These findings provide a foundation for follow-up studies aimed at mechanistic validation, biomarker development, and translational prediction of RITE risk. Collectively, this work offers both a refined gene signature and a methodological framework for dissecting the molecular determinants of venous invasion in RCC.

## Acknowledgments

This work was sponsored by the Littenberg Foundation. The funders had no role in study design, data collection and analysis, decision to publish, or preparation of the manuscript.

## Author contributions statement

Study concept and design: Ro.R., C.M. Acquisition of data: Ro.R., H.W., W.H.C., D.K. Analysis and interpretation of data: C.M., Ro.R., Ra.R. Drafting of the manuscript: C.M., Ro.R., Ra.R. Critical revision of the manuscript for important intellectual content: Ro.R. Statistical analysis: C.M. Supervision: Ro.R.

## Additional information

## Data availability

The original contributions presented in this study will be made publicly available upon publication. Transcriptome sequencing data generated for this study will be deposited in the NCBI SRA; the corresponding accession number(s) will be provided upon release. In addition, publicly available datasets were analyzed in this study. The TCGA-KIRC dataset can be found here: [https://portal.gdc.cancer.gov/projects/TCGA-KIRC], and Wang’s external dataset can be found here: [https://www.ncbi.nlm.nih.gov/bioproject/PRJNA596359 and https://www.ncbi.nlm.nih.gov/bioproject/PRJNA596338]

## Competing interests

The authors declare no competing interests.

## Notes

https://portal.gdc.cancer.gov/projects/TCGA-KIRC

https://www.ncbi.nlm.nih.gov/bioproject/PRJNA596359

https://www.ncbi.nlm.nih.gov/bioproject/PRJNA596338

### Competing Interest Statement

The authors have declared no competing interest.

